# Development of a Human iPSC-Derived “Corticospinal Tract-on-a-Chip” for Neurodegenerative Disease Research

**DOI:** 10.1101/2025.09.24.678092

**Authors:** Andriana Charalampopoulou, Arens Taga, Khalil Rust, Evelyn Luciani, Katherine Marshall, Elliot Montgomery, Anuradha Mansinghka, Richa Singh, Yang Zhao, Christine O’Keefe, Tza-Huei Wang, Arun Venkatesan, Christa W. Habela, Nicholas J. Maragakis

**Author notes:** **Corresponding Author:** Nicholas J. Maragakis, M.D. Johns Hopkins University Department of Neurology, The John G. Rangos Sr. Bldg., 855 North Wolfe Street Room 248, 2nd Floor Baltimore, MD, 21205 USA.

## Abstract

Degeneration of the corticospinal tract is a central feature in a number of neurodegenerative disorders and leads to significant disability. However, modeling corticospinal neuron (CSN) pathology and corticospinal connectivity in neurological disorders is particularly challenging. While rodent models are important for understanding early degeneration of CSN, interspecies differences in corticospinal connectivity and challenges of *in vivo* study suggest that human *in vitro* models of corticospinal biology may be ripe for development. Human induced pluripotent stem cells (hiPSC) are promising tools for overcoming intrinsic limitations that arise from physiological differences between rodents and humans. We have developed an innovative hiPSC-based microfluidic platform for modeling human CSN and spinal motor neuron (SpMN) connectivity. The incorporation of regionally specific astrocyte subtypes (cortical and spinal) in addition to CSNs and SpMNs in this newly designed system allows for the modeling of both regional and neural cell-subtype interactions. Using this model, multielectrode array electrophysiology reveals the maturation of both cortical and spinal motor neurons over the time course of 12 weeks. Retrograde labeling methods demonstrate synaptic connectivity between corticospinal and spinal motor neurons. Optogenetic strategies to selectively activate excitatory CNs attenuated by glutamate receptor antagonism confirms the functional relevance of the model. Incorporating morphological, electrophysiological and physiological measures of corticospinal connectivity, this platform is a versatile model for use in neurodegenerative disease research and for the future development of targeted CSN therapies.

**Significance Statement:** Degeneration of the corticospinal tract is a key feature of numerous neurodegenerative diseases, yet current *in vitro* models lack the anatomical and functional fidelity to study this system. We developed a human iPSC-derived “Corticospinal Tract-on-a-Chip” using a multielectrode array platform that incorporates regionally patterned cortical and spinal neurons and astrocytes. This model demonstrates structural and functional synaptic connectivity and enables longitudinal electrophysiological recordings. Critically, it supports compartment-specific manipulation and real-time analysis of CST network dynamics, capabilities lacking in existing systems. By mimicking human corticospinal physiology *in vitro*, this platform offers a novel tool for mechanistic investigation and preclinical testing of CST-targeted therapies. It holds broad relevance for studying disorders such as ALS, hereditary spastic paraplegia, and primary lateral sclerosis.

## Introduction

Corticospinal tract (CST) dysfunction is a hallmark of several neurodegenerative disorders, including amyotrophic lateral sclerosis (ALS), primary lateral sclerosis (PLS), primary progressive multiple sclerosis (PPMS), and hereditary spastic paraplegia (HSP) (1). The pathophysiology of these conditions is marked by the degeneration of long-projecting cortical neurons, or “dying-back” axonal degeneration, ultimately leading to secondary neuronal cell death. Additionally, the corticospinal tract has been hypothesized to serve as a conduit for disease propagation in neurodegenerative disorders, driven by prion-like protein aggregation and functional disruptions through neuronal hyper- or hypo-excitability (2, 3). This is prototypically exemplified in ALS, where motor neuron degeneration is hypothesized to begin focally in either the cortex or spinal cord and spreads both locally, affecting adjacent motor neurons, as well as through the corticospinal tract, affecting distant cortical and spinal motor neuron populations (3). One proposed mechanism underlying this spread is the “corticofugal hypothesis,” which posits a directional propagation of pathology from cortical upper motor neurons to their downstream targets in the brainstem and spinal cord via the corticospinal tract. However, evidence also suggests the potential for bidirectional and multisite onset, indicating a more complex pattern of disease dissemination.

Modeling the corticospinal tract in normal conditions in animal models presents significant challenges due to species-specific differences in the organization and connectivity of descending corticospinal axons (4). For example, in rodents, the majority of corticospinal projections terminate on spinal interneurons, with minimal direct input to alpha motor neurons. In contrast, primates, and especially humans, exhibit a greater proportion of corticospinal axons that form direct monosynaptic connections with spinal motor neurons, most notably those innervating distal limb muscles responsible for fine motor function (5). This includes projections from large layer V pyramidal neurons (or Betz cells) in the motor cortex, which are uniquely prominent in humans. Under disease conditions, traditional animal models of disorders involving corticospinal dysfunction have historically been utilized to provide insights into potential pathogenic mechanisms (6). However, these models also introduce significant interspecies differences that may limit their translational relevance, particularly for therapuetic applications (7). In ALS for example, mouse models have been instrumental in demonstrating early degeneration of corticospinal neurons (CSNs), supporting the hypothesis of corticofugal disease propagation (8, 9) and exploring neuroprotective strategies to promote CSN survival (10–13). However, species-specific differences in the connectivity between CSNs and SpMNs may affect the extent of CSN-related pathology in these models (4, 14–16). In diseases that affect both ends of the corticospinal pathway—such as ALS, where dysfunction of both CSNs and SpMNs is evident—the relative clinical and electrophysiological accessibility of assessing SpMN dysfunction, combined with its prominent involvement, has historically biased animal models toward a focus on SpMNs (17). Consequently, interventions that appeared effective or promising in preclinical animal models have systematically failed to demonstrate efficacy in human clinical trials, highlighting a persistent and frustrating barrier to successful therapeutic translation (18).

The use of human-induced pluripotent stem cells (hiPSCs) has emerged as a promising approach to overcome the limitations of animal models, including the ability to study diseases that lack specific gene mutations as causative events. Furthermore, to approximate central nervous system modeling, the incorporation of other non-neuronal cell types into culture systems, whether in 3D organoids or 2D co-cultures (19), has demonstrated significant enhancements in neuronal maturation, complexity, and network formation (20), increasing the physiological relevance of these models. In translational studies, co-culturing neurons with glial cells has enabled the modeling of key pathological features, such as TDP-43 mislocalization and axonal degeneration observed in ALS, HSP, and MS (21–24). Two dimensional co-cultures, in particular, offer advantages in terms of controlled culture conditions and cellular composition, enabling the integration of patient-derived cells and the evaluation of cell-autonomous or non-cell autonomous roles of specific cell types (25, 26). These systems have been effectively utilized to model local cortical and spinal cord networks, as well as motor neuron-muscle interactions (27–29).

Despite these advancements, the development of *in vitro* models to study the corticospinal tract remains largely underexplored. Previously reported systems have lacked anatomical relevance, and the versatility needed for reliable translational outcomes (27, 30, 31). To address these limitations, we have developed a novel corticospinal tract-on-a-chip model that offers a unique set of advantages. The platform uses microfluidic chambers to spatially isolate hiPSC-derived cortical and spinal neurons and astrocytes to generate region specific co-cultures. This platform mimics the anatomical organization of the human corticospinal tract, enabling the study of region-specific neuronal and astrocyte interactions as well as synaptic transmission. The microfluidic separation of individual cultures and corticospinal axons enables selective manipulation of each compartment. By incorporating optogenetic strategies for activating specific neuronal subpopulations and electrophysiological analyses with multielectrode array (MEA), this approach allows for the assessment of the dynamic properties of synaptic connectivity and network activity. This platform establishes a novel experimental framework for modeling neurodegenerative diseases affecting the corticospinal tract and enables therapeutic discovery efforts targeting corticospinal pathology.

## Results

### Generation of regionally specific hiPSC-derived neuron and astrocyte populations

We sought to design a platform that incorporated regionally-specific cell lineages, neurons and astrocytes for our purposes, into a versatile culture system that can be readily manipulated.. We differentiated healthy control human induced pluripotent stem cells (hiPSC) into cortical neurons (CN), including a subset of long projecting corticospinal motor neurons (CSN), spinal motor neurons (SpMN) as well as cortical astrocytes (CA) and spinal astrocytes (SpA) (**Figure 1A-D**). In order to maintain consistency across the experimental design, hiPSC-CN as well as hiPSC-SpMN were all derived from a single control hiPSC line (CS9XH7, Cedars Sinai). Healthy hiPSC cortical and spinal astrocytes, derived from a separate control hiPSC line (WC-30), and previously characterized for their regional identity (BX-0600 and BX-0650 respectively, BrainXell, Madison, WI, USA), were also incorporated (**Figure 1B, D**).

**Figure 1.**
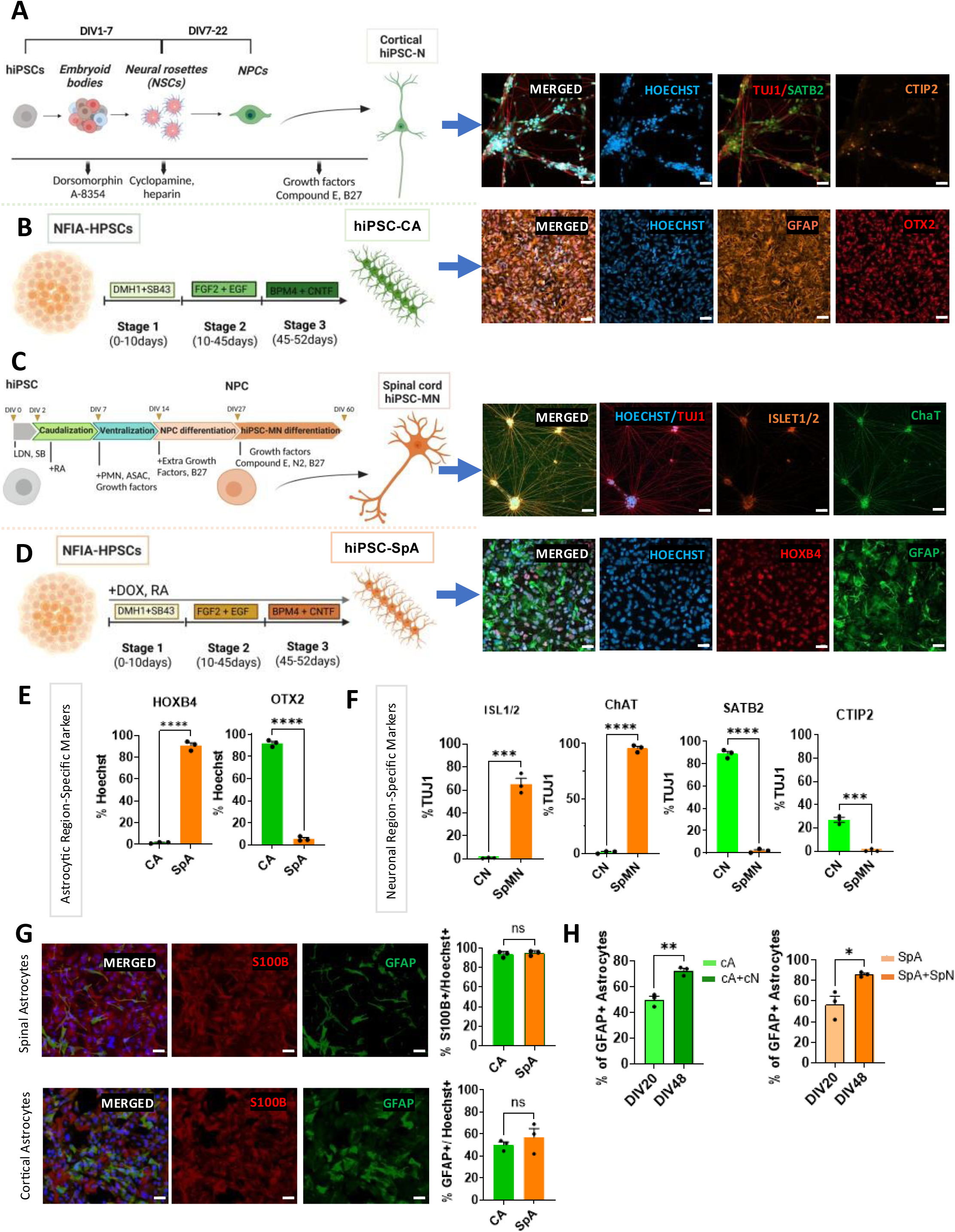
Differentiation protocols yield regional-specific cortical and spinal cord neurons and astrocytes. **A.** Schematic diagram for generation of cortical neurons from hiPSC and **B.** the diagram of hiPSC-derived cortical astrocyte differentiation of BrainXell astrocytes (modified BrainXell© illustration) as well as representative images of immunostaining for region-specific markers. **C**. Schematic diagram for generation of SpMN from hiPSC and **D**. the diagram of hiPSC-derived spinal astrocyte differentiation of BrainXell astrocytes (modified BrainXell© illustration) with representative immunostaining images showing the region-specific markers**. E**. Quantification of region-specific astrocytic markers performing immunohistochemistry showing strong regional identity as indicated by high OTX2 expression in the cortical astrocytes and high HOXB4 expression in the spinal astrocytes. **F.** Quantification of neuronal markers shows that a large subset of TUJ1+ spinal neurons display cholinergic spinal identity (ChAT+) as well as ISL1/2 expression, while cortical neuron differentiation protocol generates SATB2+ and CTIP2+ cortical neuron subtypes. **G.** Quantification of widely used astrocytic markers for both our astrocytic types to further identify them showing high levels of GFAP and S100β. **E.** Comparison of GFAP expression in two time points, DIV20 before plating with neurons (i.e. monocultures) and DIV48 (after one month of co-culture with cortical neurons and spinal motor neurons, respectively, in the chip), shows statistically significant increase of the GFAP+ levels at both cortical and spinal cocultures. T-test, * *P*<0.05, ** P<0.01. Scale bar=40μm.

Human iPSC-CN were differentiated (**Figure 1A, F)** in high purity with 91.1±0.7% immunostaining for the upper-layer cortical neuron marker, special AT-rich sequence-binding protein 2 (SATB2). A subset (27± 2.1%) of hiPSC-CN also expressed chicken ovalbumin upstream promoter transcription factor (COUP-TF)-interacting protein 2 (CTIP2), a marker of subcerebral projection CSN lineage (32).

Using methods to generate relatively pure SpMN (**Figure 1C, F**)(33), our differentiation protocol yielded 96±1.9% cholinergic motor neurons as defined by choline acetyltransferase (ChAT) immunostaining. A significant subset of these SpMN also expressed the motor neuron transcription factor ISL1/2 (65 ±8.6%). Confirming the region-specificity of the protocol, fewer than 2% of SpMN cultures exhibited region-specific cortical neuron phenotypes.

Previous studies have utilized specific differentiation methods to generate hiPSC-derived cortical (34) and spinal astrocytes (35, 36). The schematic for astrocytic differentiation was a modified version of the BrainXell protocol (36) (**Figure 1B, D**). We utilized these regionally specific astrocytes to allow for an added layer of regional specificity and fidelity to the platform. Human iPSC-derived cortical astrocytes expressed markers of astrocyte lineage (50±2.8% GFAP and 93.7±1.7% S100B) as well as a marker of cortical identity (OTX2 92±1.7%) **(Figure 1B, E, G)**. Human iPSC-SpA showed 57±7.9% GFAP and 95±1.4% S100β expression as well as a marker of spinal lineage HOXB4 (91±2.5%). Fewer than 2% (1.6±0.4%) of cortical astrocytes expressed HOXB4 and only 5.5±1.2% of SpA expressed OTX2 (**Figure 1D, E, G**). Together, these results validate the establishment of region-specific neuronal and astrocytic subtypes required for modeling corticospinal connectivity.

We have previously demonstrated that co-culturing astrocytes and neurons promotes reciprocal maturation without altering their regional identities. In the present study, we sought to confirm this observation within the current experimental paradigm. Comparing the GFAP expression of both types of regional astrocytes in monoculture and after one month in coculture with their respective neurons (neurons to astrocytes ratio 2:1), showed a significant increase in the expression levels in the co-culture with neurons (*P*= 0.0028 and *P=*0.0224 for the cortical and the spinal astrocytes respectively) (**Figure 1H**).

### Design of microfluidic device for corticospinal tract modeling

In order to provide spatial separation of the cortical and spinal compartments, we bioengineered a dual chamber PDMS (Polydimethylsiloxane) microfluidic device that allows for imaging of cell subtypes and incorporates a multielectrode array for electrophysiological study. Each chamber was calculated to have 3.69cm^2^ surface area which allowed for a plating density of 6-8 x 10^4^/cm^2^ cells per chamber. This density was optimized to ensure sufficient coverage of the electrodes while minimizing the total number of cells required. Neurite connectivity between the two chambers was accomplished by incorporating 100 microchannels. To minimize fluidic exchange but allow for cortical neurites to extend distally into the spinal chamber, the microchannels had a width of 8-10μm and a length of 450μm (**Figure 2A**). We specifically employed box-shaped PDMS microfluidic devices in our study due to several technical advantages. Their open-top design allows for homogeneous surface coating, ensuring consistent distribution of extracellular matrix proteins essential for cell adhesion and growth and, also, enables the direct addition of small molecules (as well as viral vectors) onto the culture surface, promoting uniform distribution across the entire cell population.

**Figure 2.**
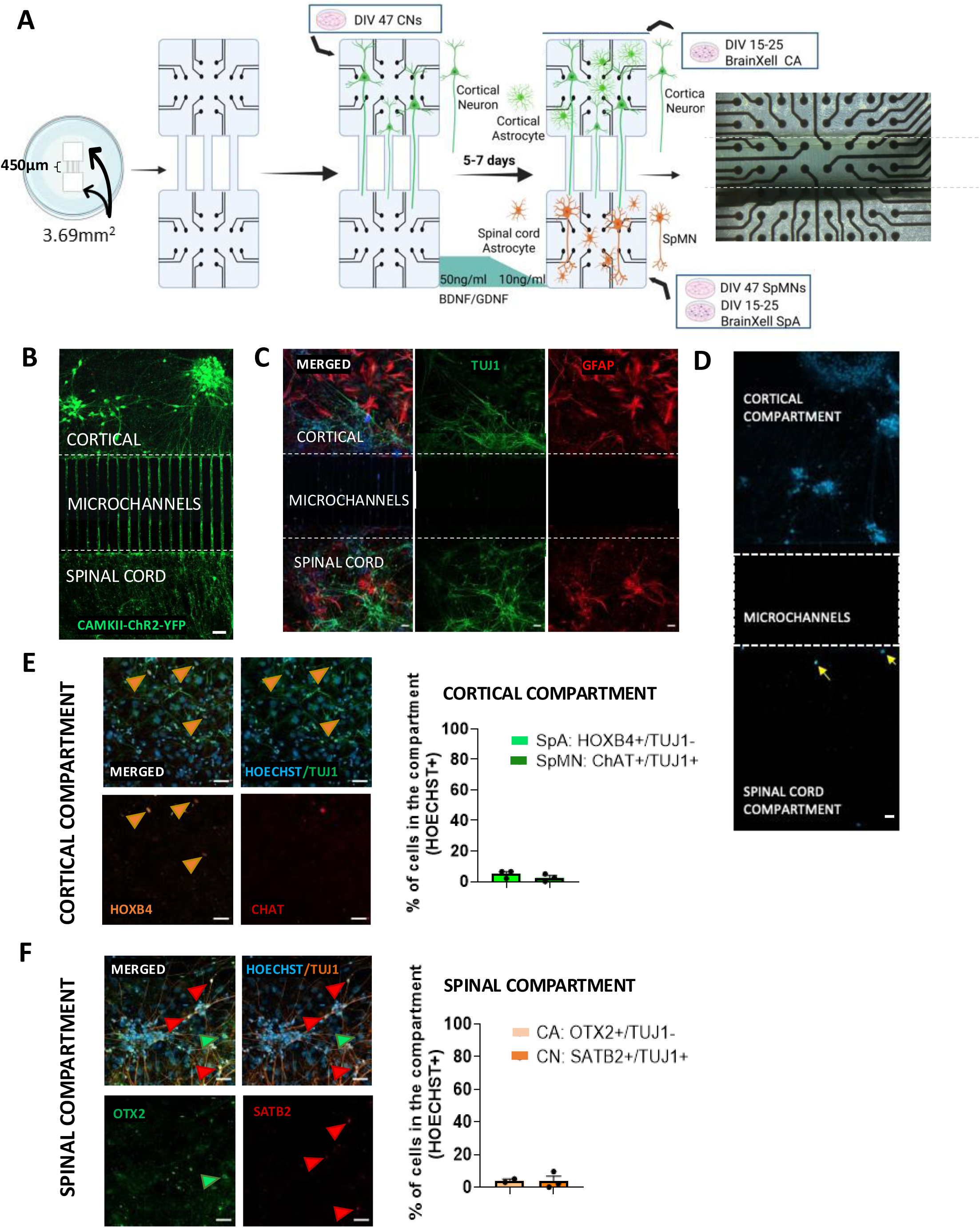
Dual microfluidic chamber design models corticospinal tract. **A.** Schematic diagram demonstrating the culture of hiPSC-CN followed by CA, SpMN and SpA on dual chambered box microfluidic devices connected through microchannels and bonded on MEA plates. **B**. CAMKII-ChR2-YFP-transduced cortical neurons identify excitatory CNs and demonstrate neurite abundant outgrowth from cortical to spinal compartment. **C**. Immunohistochemical staining identifies TUJ1+ neuron and GFAP+ astrocyte interactions in each of this dual-chambered co-culture platform. **D.** Nuclear staining (Hoechst) showing minimal migration of cells from one compartment to the other through the microchannels (8±2 cells/device, 6 devices). **E.** Quantification of cells migrating from the spinal to the cortical (orange arrows are indicative of SpA: HOXB4+TUJ1-, no ChAT+ cells) and **F.** from the cortical to the spinal compartment (green arrows indicate CA: OTX2+TUJ1-, and red arrows indicate CN: SATB2+TUJ1+), using region-specific immunohistochemical markers 1 month after seeding. Scale bars=50μm.

To encourage unidirectional cortical neurite growth from the cortical to spinal compartment, cortical neurons were seeded first, and their axons were guided through microchannels using media enriched with brain-derived neurotrophic factor (BDNF) and glial cell line-derived neurotrophic factor (GDNF) for chemoattraction to the spinal compartment. Cortical neurons were transduced with CAMKII-ChR2-YFP upon plating, enabling visualization of axon growth through the microchannels and into the spinal compartment (**Figure 2B**). After 4–6 days, CA, SpMN and SpA were introduced to complete the cortical and spinal co-cultures into a corticospinal tract-like arrangement (**Figure 2A, C**). This protocol yielded two spatially distinct microfluidically separated, compartments containing CSN and CA in one, and SpMN and SpA in the other. Significant axonal outgrowth from cortical neurons was observed within the spinal compartment (**Figure 2B**). The absence of significant media exchange between the two chambers was temporally confirmed using a live cell dye (NucSpot® Live 488). The application of live cell dye into the cortical compartment was taken up by CN within 10 minutes of application but there was no live cell staining in the spinal compartment for periods examined more than 24 hours (**Supplemental 1A-B)**.

To further confirm microfluidic separation, we evaluated whether there was significant migration of cells between the two chambers. For this purpose, cells were seeded in the cortical compartment, and migration was assessed one week later, by counting Hoechst+ nuclei in the spinal compartment, which was initially devoid of seeded cells. The quantification of Hoechst+ nuclei in this compartment confirmed minimal migration through the microchannels. On average, we observed 8.4±2 cells present in the spinal compartment per device (n = 6 devices) out of the 22,500 cells (15,000 neurons and 7,500 astrocytes) seeded at the compartment, indicating that only a small number of cells migrated to the adjacent compartment (**Figure 2D).**

Finally, to assess long-term cell migration in co-culture, reflecting conditions used in our extended experimental paradigms, we performed immunostaining four weeks after plating and assessed the presence of cortical and spinal neuronal markers. In the cortical compartment, we immunostained for spinal markers HOXB4 (from TUJ1-, SpA) and ChAT (from TUJ1+, SpMN) (**Figure 2E)**, while in the spinal compartment, we immunostained for cortical markers SATB2 (from TUJ1+, CN) and OTX2 (from TUJ1-, CA) (**Figure 2F)**. Our analysis revealed that only 4.0 ± 1.0% of astrocytes in the spinal compartment were OTX2+, and only 3.8 ± 2.9% of neurons were SATB2+ **(Figure 2F)**, suggesting their migration from the cortical compartment. Conversely, 5.0 ± 1.5% of astrocytes in the cortical compartment were HOXB4+, and 2.5 ± 1.4% of neurons were ChAT+, suggestive of a spinal origin (**Figure 2E**). These results indicate very little cellular migration between the two chambers, representing only a small fraction of the total seeded cells per compartment (15,000 neurons and 7,500 astrocytes), and highlight that the vast majority of cells remain within their designated compartments, even after prolonged culture time. It is worth noting that a small percentage of the neurons and astrocytes in monoculture (**Figure 1E-F**) also exhibit low levels of markers not typically associated with their expected lineage (“opposite” markers) suggesting that potential migration might be even more limited.

### Recapitulating the human corticospinal tract: region-specific marker expression in hiPSC-derived cultures

To confirm that the regional identity of cell subtypes was maintained following co-culture in this microfluidic platform, we examined relevant cell-subtype markers of neuronal and glial identity. After 1 month in co-culture, immunostaining confirmed the preservation of region-specific marker expression in the compartmentalized system, closely mirroring the observations in monocultures. In the cortical compartment (**Figure 3A**), 98.6 ± 0.6% of neurons expressed the upper layer neuron marker SATB2, while 27.2 ± 1.7% also expressed CTIP2, consistent with layer V CSN identity. Most cortical neurons were excitatory (CAMKII+ 93.6 ± 0.9%), with a smaller population of inhibitory neurons (GAD67+ 15.9 ± 1.8%). Cortical astrocytes retained their region-specific identity, with 94.3 ±0.7% expressing OTX2 and 72.5 ± 5.8% expressing GFAP, indicating preserved maturation in co-culture. In the spinal compartment (**Figure 3B**), neurons expressed 95.9 ± 0.7% ChAT and 87.5 ± 1.3% ISL1/2 along with 6.4 ± 1.1% GAD67, confirming their motor neuron identity and the presence of small subset of inhibitory interneurons, respectively. Spinal astrocytes showed 86.0 ± 3.1% GFAP immunostaining and 91.5 ± 0.9% HOXB4 immunostaining, supporting their astrocytic and regional identities. To evaluate motor neuron maturation in the CST platform, we characterized neurite complexity in both the cortical and spinal compartments using Sholl analysis (ImageJ Software). At 10 weeks in culture, both CN and SpMNs exhibited increased neurite complexity compared to an earlier time point (week 1), as evidenced by a greater number of neurite intersections across all radii (**Figure 3C-D**). Cortical neurons showed a peak number of intersections at a radius of 60μm, while SpMNs peaked at 40μm, indicating that, over time, cortical neurons develop more extended and elaborate arborization distally from the soma (**Figure 3C-D**).

**Figure 3.**
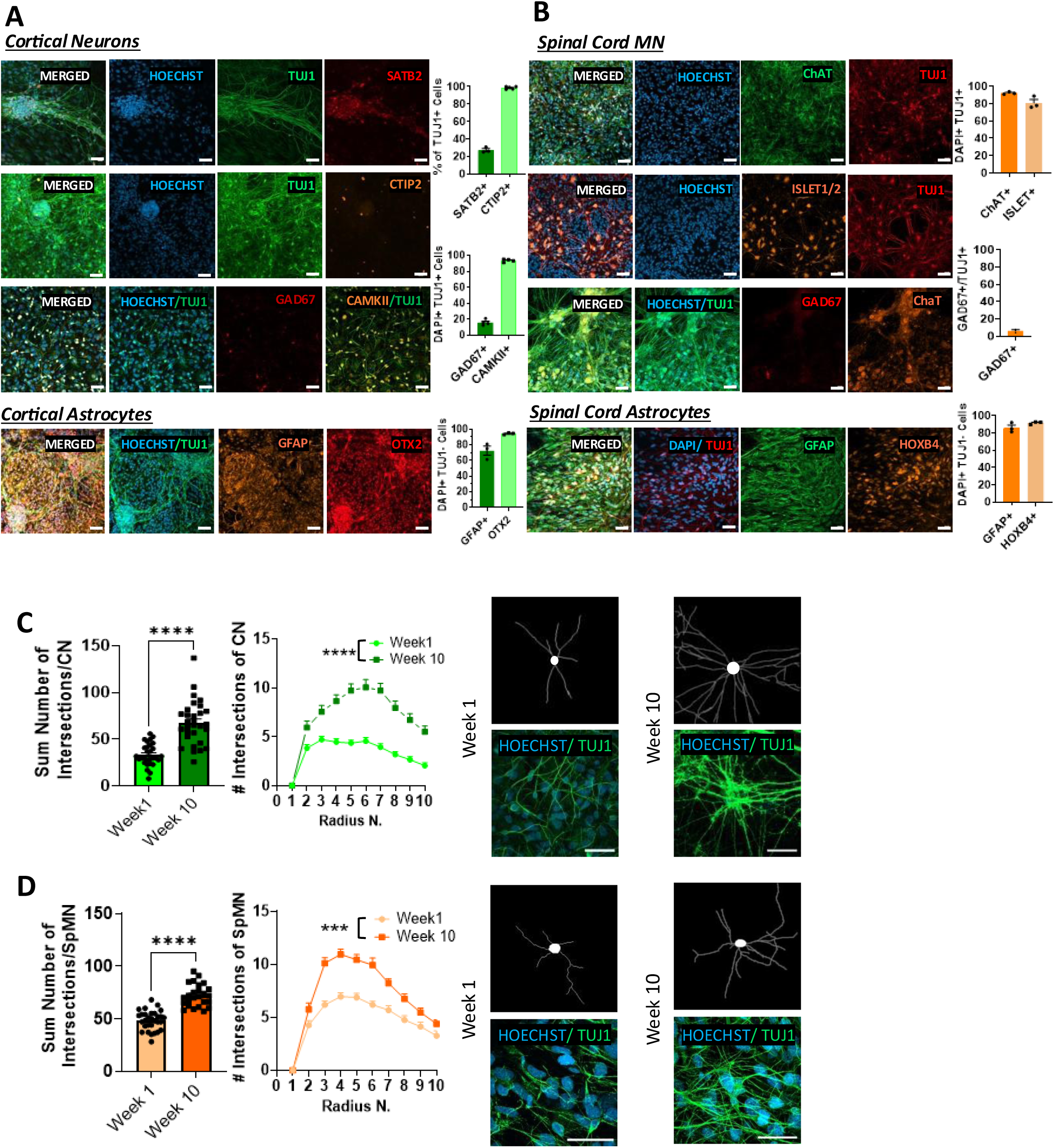
Co-culture of regional-specific neurons and astrocytes recapitulates cortical and spinal cellular maturation. **A.** Following the co-culture of CNs with CA, regional cell subtype identity is maintained with >95% SATB2 identification and 26% CTIP2+ subcerebral projection neurons. A minority of CNs expressed GAD67 indicative of inhibitory interneuron differentiation. Cortical astrocytes’ staining shows the majority of CA expressing GFAP and nearly all expressing the cortical-regional identity marker OTX2. **B**. Highly efficient SpMN differentiation was supported by elevated ChAT and ISL1/2 expression. A minority of spinal neurons expressed the inhibitory interneuron marker GAD67. Spinal astrocytes also showed high expression of GFAP and spinal-regional identity marker HOXB4. Scale bar=40μm. **C-D.** Sholl analysis of neurite outgrowth in CN and SpMN co-cultures at 1 week and 10 weeks *in vitro* (n=30 neurons per condition) with representative source and analysis images. Scale bar 50μm, One-way Anova and t-test, *** *P*<0.001, **** *P*<0.0001.

These data, taken together, demonstrate that the corticospinal tract-on-a-chip maintains region-specific identities and suggest that the morphological neuronal maturation we observed in the cocultures was not due to shifts in regional identity but was expression of emerging properties of co-cultures and their network connections through the corticospinal tract (**Figure 3A-D**).

### Electrophysiological properties and maturation of the corticospinal tract-on-a-chip

To evaluate electrophysiological maturation and emerging network activity across the corticospinal tract and within each compartment, we performed multielectrode array (MEA) voltage recordings at weekly intervals over a 12-week period. Example raster plots of spike counts per electrode demonstrated a progressive increase in activity over time (**Figure 4A**). Quantitative analysis of weighted mean firing rate (wMFR), number of active electrodes, burst percentage, average burst frequency, burst duration, and network burst frequency, revealed a consistent increase over time. Direct comparison between an early (Week1) and a late (Week 12) time point showed statistically significant increases across all parameters (*P=*0.0044, *P=*0.0001, *P=*0.0002, *P<*0.0001, *P=*0.0002 respectively) except network burst frequency that reached significancy when comparing to week 11 (*P=*0.0303) but not between week 1 and week 12. Synchrony parameters, such as the Synchrony Index and area-under-normalized-cross-correlation (AUNCC), similarly demonstrated significant increases (Week1 versus Week12, *P=*0.0036 and *P=*0003 respectively) indicating enhanced network activity over the time course of the study (**Figure 4B**). All measured electrophysiological parameters plateaued between weeks 10 and 12, suggesting the network reached a stable state of electrophysiological maturation. These results demonstrate the temporal progression of functional neuronal maturation within the model and suggest potential for studying the dynamic functional changes in the corticospinal tract.

**Figure 4.**
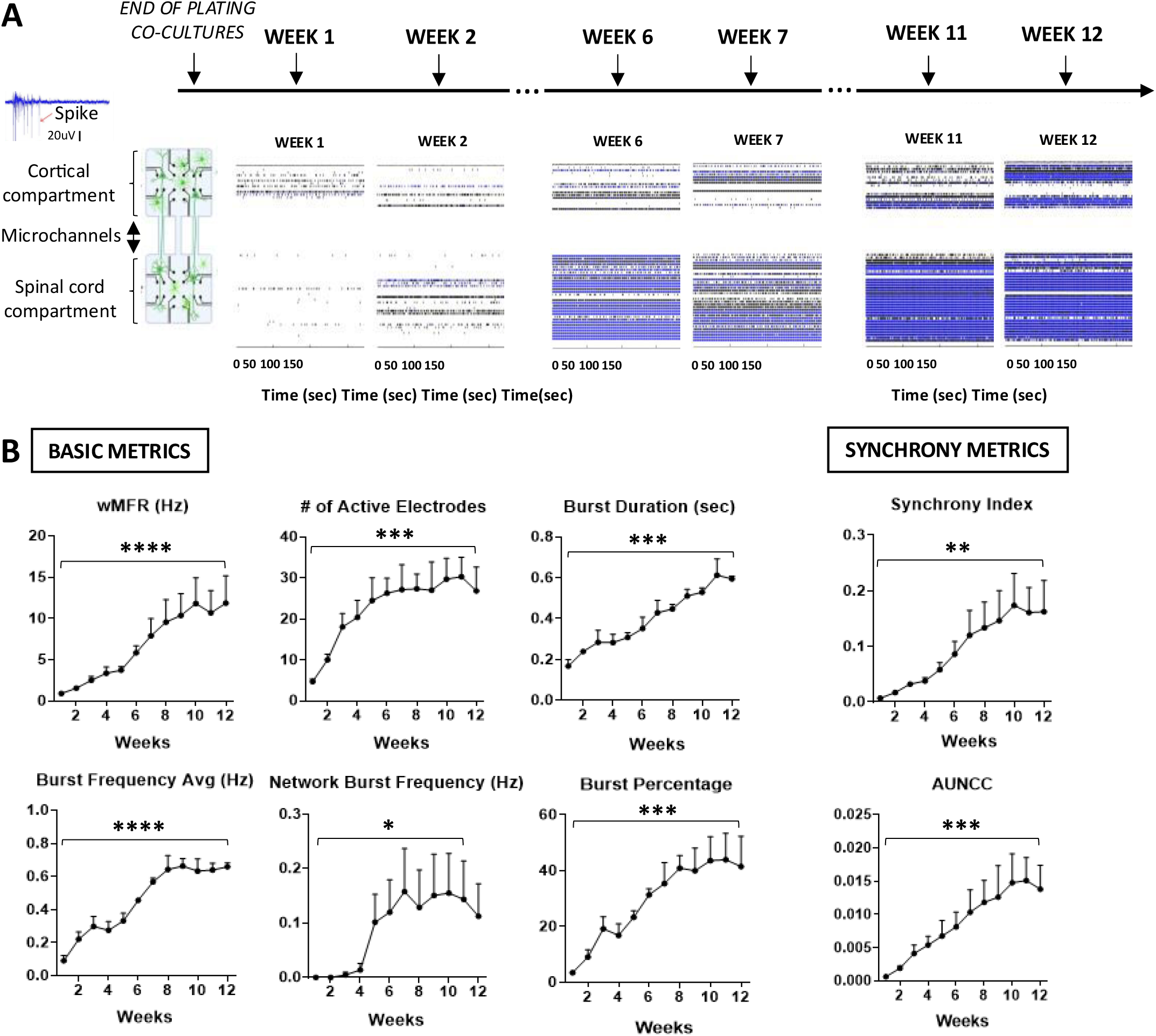
Longitudinal recordings reveal electrophysiological maturation of region-specific neuronal activity. **A**. Raster plots of longitudinal multielectrode array (MEA) recordings show increases of spontaneous activity in both the cortical (upper) and spinal (lower) compartments over 12 weeks in co-culture (microchannels between cortical and spinal compartment excluded from analysis). **B**. Quantification of electrophysiological parameters from CST-on-a-chip show increases in all basic metrics with plateaus in activity occurring in the 8-10-week co-culture time period (n=14 devices, 3 biological replicates). Comparisons refer to week 1 vs. week 12. One-way Anova, * P<0.05, **P<0.01, *** P<0.001, ****P<0.0001, wMFR: weighted mean firing rate, AUNCC: area under normalized cross-correlation.

In addition, we investigated differences in network maturation between corticospinal chips (cortical neurons and astrocytes on one side and spinal neurons and astrocytes to the other), compared to single population chips containing only a single region-specific co-culture (spinal co-cultures of neurons and astrocytes or cortical co-cultures of neurons and astrocytes on one side of the “single population chips”) (Schematic of the comparison in **Supplemental 2A-B**). MEA recordings were obtained weekly over a 12-week period to assess electrophysiologic network maturation (**Supplemental 2A-B**).

Culturing of hiPSC-CNs in cortical compartment with hiPSC-SpMNs in spinal compartment of “corticospinal chips” compared to CNs in a “single population chip” improved the average burst frequency (*P*=0.0169) as well as the burst percentage (*P*=0.045) across the time of the study (**Supplemental 2A**). Further, these hiPSC-CNs plated in corticospinal chips tended towards lower synchrony across the 12-week duration compared to CNs cultured in “single population chips”. Conversely, culture conditions in the chip did not impact spinal neuronal maturation over a 12-week time course as seen across basic and synchrony metrics (**Supplemental 2B**). These findings suggest that the presence of both cortical and spinal neuronal populations, along with their respective astrocytes, may enhance network complexity and contribute to more robust and coordinated electrophysiological activity compared to single-population systems.

### Structural evidence of synaptic connectivity between hiPSC-derived cortical and the spinal motor neuron populations

In order for our platform to model the corticospinal tract, a critical requirement was that it facilitated synaptic connections between CN and SpMN. To confirm synaptic connectivity, we employed a Rabies Virus trans-synaptic tracing strategy (**Figure 5A**). Human iPSC-derived SpMNs were infected with a G-deleted Cre-EGFP Rabies Virus (37). One week later, the helper virus AAV-CAG-FLEX-oG-WPRE-SV40-PA was introduced, enabling retrograde trans-synaptic transfer of Rabies-GFP. Cortical neurons labeled with Rabies-GFP (**Figures 5B, C**) confirmed that SpMN established trans-synaptic connections with CNs. To assess rabies-mediated retrograde transport in neurons, CNs were seeded in the cortical compartment and SpMNs in the spinal compartment, without the inclusion of their respective astrocytes. Using fluorescence microscopy to estimate the proportion of CNs connected to spinal neurons within the microfluidic device, we quantified Rabies-GFP+ CNs following trans-synaptic labeling from Hoechst+ cells (CNs). This analysis revealed that 27 ± 5.3% of CNs were Rabies-GFP+ and suggests that only a subset of CNs contributed to the corticospinal tract (**Figure 5C**). In parallel, we transduced the CN with AAV9 CAMKII-DIO-mCherry. This construct was designed to express mCherry under the CAMKII promoter and in the presence of Cre recombinase, which was provided by the Rabies GΔ Cre-EGFP virus (38). Neurons that were double-positive (GFP+ and mCherry+) were identified as synaptically-connected excitatory CNs, confirming corticospinal connectivity in this model (**Figure 5D**).

**Figure 5.**
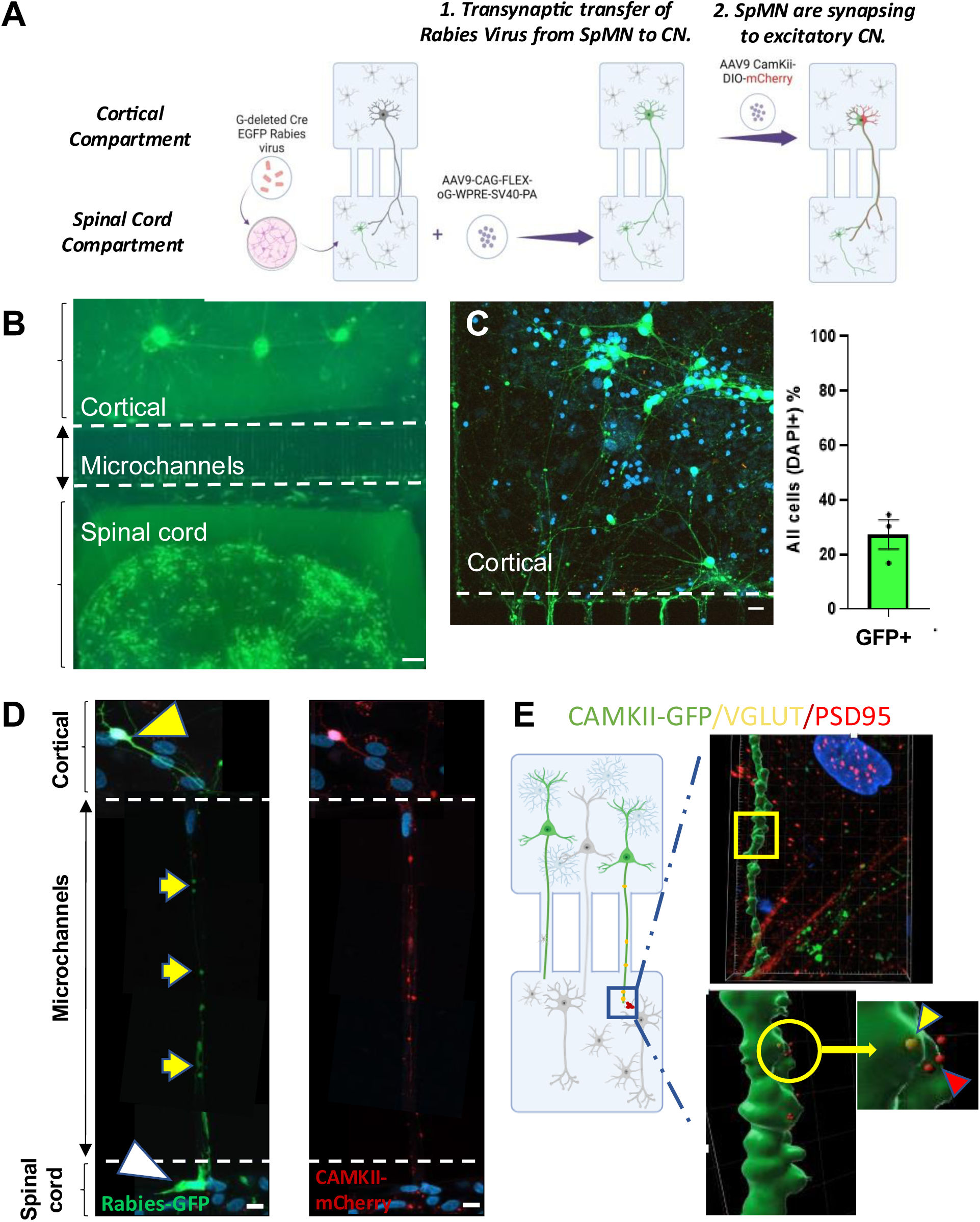
Morphologic characterization of corticospinal and spinal motor neuron connectivity. **A.** Schematic diagram demonstrating administration of G-deleted Cre-EGFP Rabies Virus to SpMNs. This was followed a week later by application of the helper virus AAV-CAG-FLEX-oG-WPRE-SV40-PA to SpMNs. AAV9 CAMKII-DIO-mCherry was added to the cortical compartment to transduce and selectively distinguish the excitatory CNs co-infected with rabies. **B.** GFP+ neurons were noted in both the spinal and subsequently in the cortical compartments indicating transynaptic viral transport from SpMN to CNs. Scale bar 100μm. **C**. Quantification of GFP+ cortical neurons one week after G-deleted Cre-EGFP Rabies Virus infection of SpMN reveals that 27.26%±5.34 CNs were synaptically connected with SpMNs. Scale bar=20μm. **D.** mCherry and GFP co-localization demonstrates the connectivity of excitatory CN subpopulations with spinal motor neurons. Scale bar=10μm. **E.** Immunostaining and subsequent Imaris 3D processing of CAMKII-GFP+ CN neurites in the spinal compartment shows co-localization (within 1µm from the axonal volume) of presynaptic (VGLUT+, minimum diameter of 0.5µm) glutamatergic marker from CN with the postsynaptic (PSD95+, minimum diameter of 0.3µm) marker from SpMN.

To further evaluate this synaptic connectivity morphologically, CNs were transduced with CAMKII-ChR2-YFP AAV, enabling visualization of their cell bodies and axons via YFP expression. We utilized Imaris software (Oxford Instruments, Oxon, UK) to generate 3D reconstructions of YFP-positive axons and qualitatively assess the spatial relationships between synaptic markers. In the spinal compartment, immunostaining with anti-YFP identified cortical axons, while antibodies against the presynaptic vesicular glutamate transporter (VGLUT) and the postsynaptic marker postsynaptic density 95 (PSD95) labeled synaptic elements. VGLUT-positive puncta (with a minimum diameter of 0.5µm) located within the surface of YFP-positive axons were considered indicative of presynaptic puncta. PSD95-positive puncta (with a minimum diameter of 0.3µm) located within 1µm from the axonal volume were interpreted as postsynaptic sites. The observed spatial proximity of VGLUT and PSD95 puncta in 3D reconstructions supported the presence of synaptic connectivity between CNs and SpMNs (**Figure 5E**).

Together, these findings provide structural evidence of synaptic connections between CNs and SpMNs within the platform, recapitulating the *in vivo* organization of long-range corticospinal projections in which cortical neurons form synapses with spinal motor neurons.

### Optical Stimulation reveals functional evidence of synaptic connectivity in the corticospinal tract-on-a-chip

Functional connectivity between cortical and spinal neurons was examined using multielectrode array recordings and optogenetic stimulation of CNs transduced with CAMKII-ChR2-YFP. Immunostaining of transduced neurons confirmed co-localization of CAMKII-ChR2-YFP with the subcerebral projection neuron marker CTIP2 and the pan-neuronal marker TUJ1, indicating the expression of ChR2 in excitatory CSNs (**Figure 6A**). Corticospinal tract-on-a-chip cultures were matured on MEA plates and stimulated with blue light (470nm) while recording the response from cortical and spinal compartments to determine light-evoked neuronal activity and synaptic transmission through the corticospinal network. Peristimulus spike count normalized to baseline activity demonstrated an immediate increase in spiking in the cortical compartment, followed by a delayed response in the spinal compartment, indicative of a temporal lag from synaptic transmission (**Figure 6B)**.

**Figure 6.**
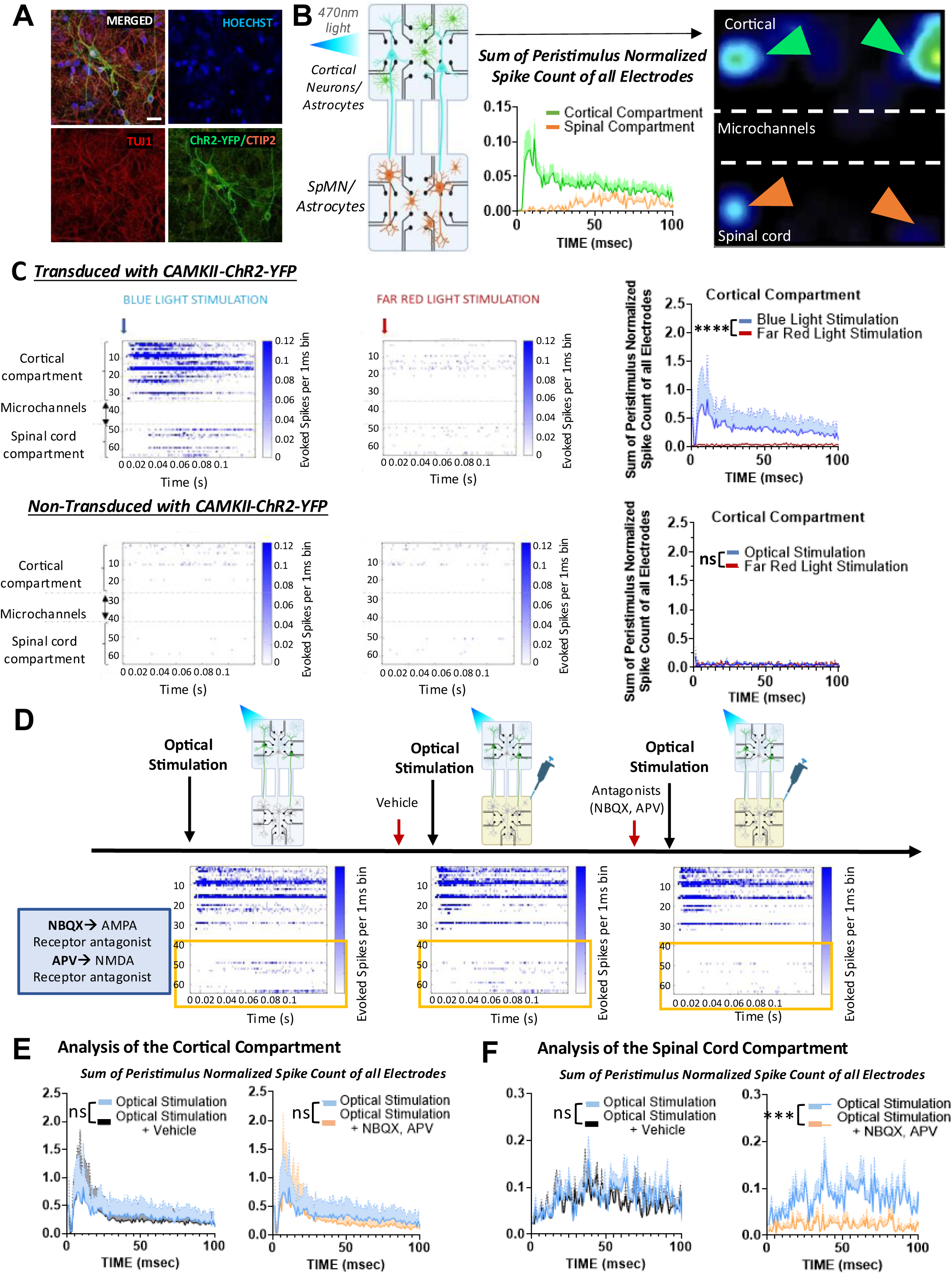
Optogenetic stimulation demonstrates network connectivity between excitatory CNs and SpMNs. **A**. CAMKII-ChR2-YFP+ transduced in CN co-localizes with subcerebral CTIP2+ in subcerebral projection CSNs. Scale bar=20μm. **B**. Optic stimulation of CAMKII-ChR2-YFP+ CN induces synchronized spike activity in MEA electrodes. Following optic stimulation, cortical neurons show summation of electrophysiological activity (green). This is temporally followed by electrophysiological responses from SpMN in the spinal compartment (orange). Visual heatmap representation demonstrates this synchronized activity across the CST. **C**. Peristimulus raster plots (1 msec bin) of CAMKII-ChR2-YFP+ transduced CN show activity in the cortical compartment followed by an electrophysiological response in the spinal compartment (top, left). Far red-light stimulation does not activate these CNs (top, right). Similarly, blue light does not activate non-transduced CNs nor does far red-light stimulation (bottom). Peristimulus plots are extracted from 5-minute blue light and far-red light stimulation experiments (70 light pulses) of 10-week co-cultures (N=3). Wilcoxon matched-pairs signed rank test **** P<0.0001. Error bars ± SEM are presented as shaded blue area. **D.** Peristimulus raster plots (1 msec bin) recorded following the addition of the AMPA receptor antagonist NBQX and NMDA receptor antagonist APV to the spinal compartment shows the reduction in electrophysiological activity in the spinal compartment with sparing of activity in the cortical compartment. **E**. Peristimulus electrophysiological activity in the cortical compartment remains unaffected regardless of the treatment conditions (N=4). **F.** Glutamatergic activity is reduced in the spinal compartment following application of glutamate antagonists (N=4). Stimulation parameters: 70 stimuli, 0.25 Hz, 5msec stimulus duration, 30% intensity. Error bars are represented as shaded area and indicate mean ± SEM. One-way Anova *** *P* <0.001, \*\*\*\**P*<0.0001.

To confirm that the observed neuronal response was specifically mediated by ChR2-YFP activation, we repeated the experiment comparing blue to far-red light stimulation (**Figure 6C**). While there was robust activation of CNs in response to blue light, far-red light elicited no significant response (P <0.0001) (**Figure 6C**). To further validate the specificity of ChR2-YFP–mediated activation, blue light stimulation of non-transduced hiPSC-CNs (i.e. CNs not expressing ChR2-YFP) did not elicit any electrophysiological response, with no significant difference compared to far-red light stimulation (P = 0.9493; **Figure 6C**). Peristimulus spike count showed that blue light stimulation of CNs induced a distal response in SpMNs (**Figure 6F**). These findings demonstrate that optical stimulation of CNs induces a synaptic response in SpMNs, suggesting functional connectivity between the two neuronal populations in this platform through the corticospinal tract axons.

To investigate whether the SpMN response to cortical optical stimulation was mediated by the neurotransmitter glutamate, we utilized a pharmacological approach and considered three conditions: (1) blue light stimulation under baseline conditions, (2) blue light stimulation following the addition of vehicle solution, and (3) blue light stimulation following the addition of a cocktail containing the AMPA receptor antagonist 2,3-dihydroxy-6-nitro-7-sulfamoyl-benzo[f]quinoxaline (NBQX) and the NMDA receptor antagonist 2-Amino-5-phosphonovaleric acid (APV) (**Figure 6D**). Raster plots showed a consistent electrophysiological response from hiPSC-SpMN following the optical activation of CAMKII-ChR2-YFP+ hiPSC-CNs under baseline and vehicle conditions (n=4 biological replicates). Quantification revealed no statistical difference in either compartment in the peristimulus spike count before and after the addition of vehicle (**Figure 6E**). However, blocking AMPA and NMDA receptors with NBQX and APV led to a significant decrease (*P=*0.0009) in spinal compartment activity **(Figure 6F**), while hiPSC-CN activity remained unaffected (**Figure 6E**). These findings confirm that the spinal neuronal response to cortical optical stimulation is mediated by glutamatergic transmission, providing further evidence of functional synaptic connectivity within the corticospinal tract-on-a-chip.

## Discussion

Modeling corticospinal connectivity in healthy control conditions and in neurological disorders presents a distinct challenge due to the inherent structural and functional complexity of this long-range projection pathway. Traditionally, studies of the corticospinal tract and its synaptic connectivity have relied heavily on animal models (39) with notable limitations in capturing the cellular and molecular mechanisms underlying human pathobiology (40, 41). Non-human primate models may provide a closer approximation to human neuroanatomy and disease mechanisms but are also limited in their ability to recapitulate neurodegenerative disorders that involve complex or unidentified genetic and environmental etiologies, such as PPMS and sporadic ALS. In addition, their use is constrained by significant ethical, logistical, and financial challenges (42, 43). The development of a humanized corticospinal-tract-on-a-chip is our innovative approach to address some of these challenges.

Human iPSC-derived neuronal cultures are emerging methods for examining a host of intrinsic cell properties in both health and in neurodegenerative disease (44–49). Donor-derived iPSCs enable the generation of disease-relevant neuronal subtypes, including corticospinal neurons and spinal motor neurons, which have the potential to recapitulate key pathological features of disorders with prominent corticospinal pathology, including ALS and HSP, such as TDP-43 mislocalization and axonal degeneration (23, 44). For this platform, we employed optimized protocols to generate highly pure populations of cortical neurons, spinal motor neurons and already highly characterized region-specific astrocytes from human iPSCs. Our findings notably demonstrate that the iPSC-derived cell subtypes acquire molecular signatures characteristic of their *in vivo* counterparts. Approximately one-fourth of the cortical neurons exhibited characteristic layer V markers such as CTIP2 that has been described *in vivo* as indicative of subcerebral projection neurons shown to degenerate in models of neurodegeneration (10). Moreover, spinal neurons expressed ChAT and ISL1/2 in high abundancy, while astrocytes were regionally patterned to express forebrain (OTX2) or spinal (HOXB4) transcription factors, as astrocyte heterogeneity is recognized as important to both normative biology and pathology (50–52). *In vivo*, astrocytes from different brain regions such as the cortex, hippocampus, midbrain, and cerebellum, exhibit distinct synaptogenic profiles due to variations in gene expression of synaptogenic factors (53, 54).

One of the goals of this study was to engineer a chip with a more physiologically relevant microenvironment. With these four fully characterized cell types, we aimed to bridge this gap combining these populations in an integrated manner, while most studies using human iPSC-derived CSNs and SpMNs have utilized monocultures or simple co-cultures with glial cells. It is well established that co-culturing neurons with astrocytes enhances synaptogenesis, with astrocyte-conditioned media alone sufficient to increase synaptic density (55). Additionally, direct neuron-astrocyte interactions influence astrocyte gene expression (56) as also evidenced by increased GFAP expression in astrocytes co-cultured with neurons compared to monocultures (19). In this study, we showed that co-culturing neurons and astrocytes in our system also resulted in increased GFAP expression compared to astrocyte monocultures, further supporting the role of neuronal interactions in astrocyte maturation and vice versa (57, 58). Future studies could incorporate additional cell types, such as oligodendrocytes and microglia, to further enhance the physiological relevance of the model. This expansion may provide deeper insights into cell-type-specific contributions and interactions within the corticospinal circuit, building upon the foundational framework established with the current two-cell-type system.

To combine the cell types in a single design, we leveraged recent advances in bioengineering technologies that have expanded the capabilities of iPSC-based platforms. Some platforms enable the generation of 3D brain and spinal cord organoids that self-organize to mimic aspects of human neural development, including cortical-cortical and motor unit connectivity, and synapse formation (59, 60). Building on these developments, the integration of microfluidic devices and scaffold-based assembloids has further enhanced the ability to interrogate synaptic biology, axon guidance, and long-range connectivity in a controlled and modular fashion (60, 61). These platforms have been used to model long-range projection systems, including the visual, somatosensory, and motor pathways (38, 62). Considering both the strengths and the limitations of these 3D systems (e.g. reduced experimental control over individual cell types and media conditions, limited spatial and temporal resolution, challenges in quantitatively assessing functionally and morphologically relevant outcomes), we underscore the critical need for refined platforms that combine anatomical fidelity with experimental versatility to effectively model corticospinal connectivity.

To address these requirements, we developed a human iPSC-based *in vitro* model of the corticospinal tract that integrates the scalability and accessibility of 2D cultures with the spatial resolution afforded by microfluidic compartmentalization. Simple and more sophisticated microfluidic designs, often referred to as “lab-on-a-chip” systems, have paved the way for innovative approaches to studying the pathophysiology of neurodegenerative diseases using these region-specific iPSCs (63, 64). We chose an open-top style compartmentalized box device that required small numbers of cells, as well as allowed for long-term survival and homogeneous distribution of neurons that was important for the longitudinal assessment of neuronal morphology and electrophysiology

To further replicate corticospinal tract morphology, an emphasis on unidirectional neurite outgrowth was planned. The plating of corticospinal neurons prior to SpMN as well as the encouragement of unilateral growth of neurites with the addition of growth factors helped to ensure this occurred. Filling the narrow width of the microchannels with corticospinal neurites also discouraged retrograde growth and significant cell migration across the compartments as was indicated by our results.

Importantly, we show that cortical neurons plated in this dual-chambered microfluidic system retain high cortical identity without acquiring spinal signatures. Similarly, spinal motor neurons maintained high purity in both monoculture and dual-chamber formats, confirmed by ChAT and ISL1/2 expression. We propose that this platform will be useful for modeling neurodegenerative diseases involving axonal transport deficits, synaptic dysfunction, and neuron-glia interactions, areas that are difficult to investigate using traditional culture formats (60, 61).

Our system further incorporates multielectrode array technology to enable longitudinal assessment of electrophysiological maturation and network activity of the human iPSC-derived corticospinal tract over time. By ensuring coverage of both compartments with electrodes, the platform supports simultaneous electrophysiological measurements from cortical and spinal chambers. Our data demonstrate direct measurement of neural activity at both compartments as well as progressive increases in spontaneous activity and corticospinal synchrony over time, with an electrophysiological plateau observed around 8–10 weeks of co-culture.

We and others have previously described MEA-based physiological maturation of SpMN, following culture with astrocytes (19). Studies comparing neuronal cultures originating from the prefrontal cortex, hippocampus, amygdala and spinal cord have shown unique electrophysiological patterns for each subtype. Specifically, changes in spike rate, interspike interval, time between bursts, and burst frequency were found to be region specific for human iPSC-derived neurons. While Dauth et al. reported minimal differences in electrophysiological metrics like firing rate and interburst interval across neuronal subtypes within a multiregional brain-on-a-chip model, they observed clear differences when these populations were compared to their respective monocultures (65). This indicates that the functional interaction between regionally distinct neuronal types alters their maturation dynamics and shapes the emergent electrophysiological properties of the network (66). In our analysis, we observed a trend towards lower levels of synchrony in cortical cocultures from the corticospinal chips, compared to cortical neurons from the single population chips. However, we did not find a similar effect on the electrophysiological maturation of SpMN from the corticospinal chips compared to the single population chips that had similar electrophysiological pattern of maturation in the parameters examined. It is worth noting that synchrony of SpMN in the corticospinal chips had a trend towards lower levels compared to the single population chips, showing a firing pattern closer to human physiology. These results support the idea that corticospinal interactions influence electrophysiological maturation.

Electrophysiological studies have increased the appreciation for demonstrating neuronal maturation, particularly when using *in vitro* iPSC cultures (67). Our findings highlight the importance of longitudinal time-course analysis *in vitro* in order to develop a more accurate representation of *in vivo* biology. The maturation and the stabilization of electrophysiological activity may be particularly relevant in the modeling of neurodegenerative disorders and should be taken into consideration in studies using humanized corticospinal tracts from iPSCs to investigate either health or disease states.

Our MEA data show, as mentioned before, that independent and simultaneous recordings from either the cortical or spinal compartments were possible, allowing for studies that could examine the interplay between CN and SpMNs. In addition, pharmacological blockade experiments proved that the selective inhibition of synaptic activity following the application of glutamate antagonists to the spinal compartment did not affect the cortical compartment, which also suggests that there is fluidic isolation of these compartments and the platform affords the ability to study these populations in isolation.

One more critical feature of this model is the ability to recreate appropriate corticospinal synaptic structure, synaptic receptor subtype, and synaptic transmission. In the literature, limited attention has been given to modeling the functional and structural connectivity between these neuronal populations in the context of a more complex network. To make an effective and relevant model, we wanted to demonstrate efficient connectivity between cortical neurons and SpMN. One approach to do this necessitated a retrograde tracing strategy with Rabies virus, using a modified method described before for retrograde transport and synaptic connectivity in human-derived aggregoids. The use of AAV9 CAMKII-DIO-mCherry transduction in cortical neurons also demonstrated that this connectivity was relevant to excitatory cortical neurons (36). A second confirmation was investigated using immunohistochemistry. We demonstrate that hiPSC-cortical neurons, identified by CAMKII-ChR2-YFP, express the vesicular glutamate transporter VGLUT localized to neurites within the cortical compartment as well as in distal neurites extending, through microchannels, into the spinal compartment. Importantly, the presence of VGLUT at these distal neurites was associated with synapses to cholinergic SpMN as defined by postsynaptic PSD95 immunostaining.

To firmly establish that the morphological connectivity we described was physiologically and functionally relevant, we utilized a CAMKIIα-ChR2-YFP construct to selectively transduce excitatory cortical neurons, enabling blue light–mediated activation of channelrhodopsin. This approach allowed us to synchronize cortical neuron depolarization and assess corresponding responses in SpMNs. This corticospinal activity appears to be mediated by glutamatergic excitatory neurotransmission since blocking of AMPA and NMDA receptors resulted in a suppression of the electrophysiological response in SpMN.

Taken together, our data provide a foundation for a humanized corticospinal tract-on-a-chip that includes the incorporation of region-specific neural cell subtypes and exhibits morphological and electrophysiological characteristics that could prove useful for the study of neurodegenerative diseases with prominent cortical and spinal motor neuron pathology. Ultimately, this two-dimensional corticospinal–tract–on–a–chip holds significant potential for studying corticospinal connectivity in CNS disease modeling, providing a platform for both mechanistic investigations and pharmacological studies.

## Materials and Methods

### Microfluidic Device fabrication

Microfluidic devices were fabricated using standard photolithography and soft lithography techniques as previously described (68, 69). Photomasks were designed using AutoCAD software and printed by Artnet Pro. Briefly, a silicon wafer was dehydrated by overnight baking at 200 °C. The SU-8 3010 photoresist (Kayaku) was spun on the wafer at 3000 rpm to form microchannels layer and SU-8 3050 photoresist was spun subsequently on the wafer at 1000 rpm. The wafer was baked at 95 °C for 45 mins. After exposure, the wafer was developed and baked at 200 °C for an hour. PDMS and cross-linker (Electron Microscopy Sciences, Pennsylvania) were mixed at a 10:1 ratio (w/w), degassed under vacuum, and cast onto the molds to ensure complete coverage of the microchannels. The molds were cured in an oven at 80 °C for 2 hours, after which the cured PDMS was carefully peeled off. Square compartments, open at the top to facilitate cell seeding and media addition, were created by punching out the desired regions of the PDMS. The resultant compartmentalized devices consisted of two chambers (surface area of 3.69 mm²) connected by 100 microchannels with diameters ranging from 8 to 10μm. After sterilization, the devices were bonded by plasma treatment to glass-bottom 12-well plates for imaging studies or to 6-well multielectrode array plates from AxionBiosystem for electrophysiological studies. Prior to cell plating, the surfaces of the box devices were coated with GeltreX™ (Gibco).

### Differentiating cortical NPCs from hiPSCs

Human iPSC derived cortical neural progenitor cells (NPCs) were generated from a control line (CS9XH7) using the human forebrain cortical differentiation protocol as adapted from Wen et al 2014 (34). Briefly, iPSC colonies were detached from the Matrigel® (Corning) with 1 mg/mL collagenase for 1h and suspended in embryoid body (EB) medium, consisting of DMEM/F12 (-) L-Glutamine, 20% knockout serum replacement (KSR), penicillin-streptomycin (Pen/Strep), GlutaMax, non-essential amino acids (NEAA), supplemented with [55μM] β-Mercaptoethanol (Gibco), [2μM] dorsomorphin, and [2μM] A-83, in low-attachment plates (Corning) plates for 4 days with daily medium exchanges. After 4 days *in vitro* (DIV), EB medium was replaced by NPC induction medium (“NPC medium”) consisting of DMEM/F12, N2 supplement, NEAA, 2μg/mL heparin and [2μM] cyclopamine. The floating EBs were then transferred to Matrigel-coated 6-well plates on day 7 and allowed to form neural rosettes. The attached rosettes were maintained for 15days in NPC medium, changed three times per week. After 22DIV, neural rosettes were selected and lifted mechanically after incubation with neural rosette selection reagent (StemDiff™, STEMCELL technologies) for 1 hour and transferred to low attachment plates (Corning) in NPC medium enriched with B27 supplement (Gibco). Resuspended neural progenitor spheres were dissociated with accutase (Gibco) at 37°C for 10 min and frozen in “Cortical Neuronal Medium” consisting of Neurobasal base (Gibco) with B27, NEAA, GlutaMax, 10 ng/mL BDNF, 10 ng/mL GDNF and supplemented with 10% DMSO for cryopreservation(70).

### Differentiating cortical neurons and astrocytes from cortical NPCs

After 23 DIV, cortical NPCs were thawed and plated onto plates that had been first treated with Poly-L-ornithine (PLO) diluted in water for 1 hour at 37°C and then coated with 1:100 mouse laminin (1μg/mL, Gibco). Frozen NPCs were flash thawed in a 37°C water bath, centrifuged and resuspended in 10 ml of cortical neuronal media with 20μM of ROCK-I (1:500 DMSO) and added to the laminin coated plates. For cortical neuron maturation, during the first week after plating, neuronal cultures were treated for 48 hours with 0.2μM cytosine arabinoside (ARA-C) in neuronal media to suppress astrocyte/glial proliferation. Cells were maintained with media changes three times a week from DIV23 to DIV44. The media consisted of cortical neuronal medium supplemented with [125nM] Compound E (Santa Cruz Biotechnology) to further enhance motor neuronal maturation (71).

Human iPSC-derived cortical astrocytes derived from a separate control hiPSC line (WC-30) were purchased from BrainXell and differentiated per manufacturer’s recommendation for a week. Then, cells were cultured with Astrocyte Differentiation medium with 1% fetal bovine serum (FBS) consisting of DMEM/F12, GlutaMax, NEAA, 1%FBS, Pen/Strep, B27 supplement and 2μg/ml Heparin. Cells were expanded into Matrigel-coated plates and passaged upon reaching confluency until DIV15-25 when they are used for seeding.

### Differentiating spinal NPCs from hiPSCs

Induced pluripotent stem cells from the same control line (CS9XH7) were differentiated into spinal neural progenitor cells (NPCs) following modifications of a 25-day protocol described previously (33) that relies on extrinsic morphogens to pattern NPCs along the rostro-caudal and dorso-ventral axis. Briefly, key steps include neuralization via dual SMAD (“Suppressor of Mothers against Decapentaplegic”) signaling inhibition using LDN193189 (Stemgent) and SB431542 (Millipore Sigma), followed by caudalization and ventralization through retinoic acid (RA) and purmorphamine (PMN) (Millipore Sigma).

### Differentiating spinal neurons and astrocytes from spinal NPCs

25DIV spinal NPCs were thawed and plated onto PLO and Laminin-coated plates as described above for cortical NPCs. For motor neuron differentiation, spinal NPCs were cultured with “spinal neuronal differentiation medium”, comprised of Neurobasal (Gibco), enriched with N2 and B27 supplements (Gibco), GlutMax, NEAA, pen/strep and supplemented with growth factors (RA, PMN, ascorbic acid, recombinant human-brain-derived neurotrophic factor, glial cell line-derived neurotrophic factor, insulin-like growth factor 1, ciliary neurotrophic factor). Similar to the cortical differentiation protocol, Compound E (Santa Cruz Biotechnology) was added to enhance neuronal differentiation into motor neurons. To prevent astrocyte over-proliferation, neuronal cultures were treated once between DIV32-35 with 0.02μM Ara-C for 48h. The medium was then changed every other day. At 60 DIV, this protocol has been shown to generate a population of spinal cord neurons, with a majority of neurons expressing spinal motor neuron markers, including choline acetyltransferase (ChAT) and ISL1/2 (19).

Human iPSC-derived spinal astrocytes (control line WC-30) were purchased from BrainXell and differentiated per manufacturer’s recommendation for a week. Cells were then cultured with astrocyte differentiating medium with 1% FBS as described above for cortical astrocytes.

### Generation of region-specific co-cultures

The surfaces of the chips were coated with Geltrex overnight. Cortical neurons were first seeded in the cortical compartment, we chemoattracted the axons using enriched BDNF (50ng/ml), GDNF (50ng/ml) media for 4-6 days and then the device was seeded with other cell types to complete the corticospinal-tract-on-a-chip (CA in the cortical compartment and SpMN, SpA in the spinal compartment). Cell densities in the chip were 6 x 10^4^/cm^2^ and 8 x 10^4^/cm^2^ (ratio neurons: astrocytes, 2:1), at DIV47 and DIV15-20 for neurons and BrainXell astrocytes, respectively. These densities generated a confluent layer of astrocytes with interspaced neurons, as described previously (19). In the chip, given the differing media requirements of cortical and spinal motor neuron monocultures, we developed a composite medium by combining the cortical and the spinal media recipes (72–74). Composite medium for the co-cultures in the devices included Neurobasal: DMEM:F12 at a 50:50 ratio, supplemented with B27, N2, BDNF, GDNF and ascorbic acid.

### Cell migration assessments

Cells were seeded into the cortical compartment using the standard protocol described above, and migration into the spinal compartment was evaluated after one week. Given there were no cells in the spinal compartment (opposite compartment from the one where cells were seeded) upon plating, we could identify migrated cells by counting the absolute number of Hoechst+ nuclei in the spinal compartment. Migration data was collected from a total of 6 devices and 3 field images per device were collected. To evaluate cell migration after one month in the corticospinal-tract-on-a-chip, immunostaining was used to detect cortical and spinal region-specific markers in the spinal and cortical compartments, respectively. We stained the cortical compartment for ChAT (a marker of spinal motor neurons) and HOXB4 (a marker of spinal astrocytes), and the spinal compartment for SATB2 (a marker of cortical neurons) and OTX2 (a marker of cortical astrocytes). Immunoreactive cells were quantified as a percentage of the total number of neurons (TUJ1+/Hoechst+) and astrocytes (TUJ1-/Hoechst+), based on five images per compartment from three separate chips.

### Media exchange assessment

To assess media exchange and small molecule diffusion across microchannels, we utilized 488-NucSpot 1000X (Biotium), a live nuclear fluorescent dye, as a probe to track small molecule diffusion. Six chips with all cell types seeded were used for this experiment. The live-cell dye was applied to the cortical compartments of three chips, while the remaining three received an equivalent volume of vehicle (DMSO) as control. Live imaging was performed using confocal microscopy at 5x and 10x magnification and diffusion was assessed at various time points. At the 24-hour time point, the nuclear dye was also added to the spinal compartments to verify the expected diffusion pattern. Fluorescence intensities were maintained for both compartments at each time point. Images were analyzed using Zeiss Zen software (Carl Zeiss). For representative images, adjacent tiles were acquired with overlapping regions to ensure field continuity and manually aligned into a mosaic using PowerPoint (Microsoft Office 365). Care was taken to preserve spatial continuity and anatomical context between adjacent fields. No image content was altered beyond alignment and cropping.

### Immunocytochemistry

Cells were fixed with 4% paraformaldehyde for 10 minutes and then washed with phosphate-buffered saline (PBS) three times. The cells were then permeabilized with 0.1% Triton™ X-100 (Millipore Sigma) in PBS for 10 minutes and washed with PBS three times. A blocking solution with 3% bovine serum albumin (BSA) in PBS was then applied for 1 hour. Glass bottom devices were stained with primary antibodies in a blocking solution containing 3% BSA in PBS and 3% species specific serum and incubated overnight at 4°C. The next day, cells were washed with 3% BSA in PBS three times and incubated with appropriate secondary antibodies (Thermo Fisher Scientific, Alexa Fluor Dyes; concentration: 1:1000) and Hoechst® (Thermo Fisher Scientific; concentration: 2 μg/ml) in a blocking solution with 3% BSA in PBS and 3% species-specific serum, for 1 h at room temperature. Finally, the coverslips were washed with 3% BSA in PBS three times and mounted with Prolong gold with DAPI® (Thermo Fisher Scientific) and stored at 4°C until ready to image. The primary antibodies used for this study are listed in **Tables 1-2**.

**Table 1:** Primary antibodies utilized for immunohistochemistry.

| Primary antibody | Species | Company (cat no) | Dilution |
| --- | --- | --- | --- |
| Anti-βIII tubulin | Rabbit | Millipore Sigma (AB9354) | 1:500 |
| Anti-βIII tubulin | Mouse | Abcam (AB78078) | 1:1000 |
| Anti- βIII tubulin | Chicken | Millipore Sigma AB9354 | 1:500 |
| ISL 1/2 | Mouse | Developmental Studies Hybridoma Bank (39.4D5, concentrate) | 1:50 |
| ChAT | Goat | Millipore Sigma (AB144P) | 1:100 |
| Anti-GFP | Chicken | Aves GFP-1020 | 1:1000 |
| VGLUT | Mouse | Synaptic Systems | 1:100 |
| SATB2 | Rabbit | Abcam (AB92446) | 1:100 |
| CTIP 2 | Rat | Abcam (AB18465) | 1:200 |
| GAD 67 | Mouse | Millipore Sigma (MAB5406) | 1:500 |
| CAMKIIa | Rabbit | Thermo Fisher Scientific PA5-14315 | 1:100 |
| S100B | Mouse | Millipore Sigma (ABN59) | 1:500 |
| GFAP | Chicken | Millipore sigma (AB5541) | 1:200 |
| HOXB4 | Rat | Developmental Studies Hybridoma Bank (I12, concentrate) | 1:25 |
| OTX2 | Mouse | Invitrogen (MA5-15854) | 1:400 |
| Anti- βIII tubulin (TUJ1) | Chicken | Millipore sigma (AB9354) | 1:500 |

**Table 2:** Secondary antibodies utilized for immunohistochemistry.

| Secondary antibody | Company (Cat No) | Dilution Factor |
| --- | --- | --- |
| Hoechst 33342 | Millipore Sigma 34580 | 1:1000 |
| Alexafluor™ Goat anti-chicken 488 | ThermoFisher Scientific A11039 | 1:1000 |
| Alexafluor™ Goat anti-chicken 647 | ThermoFisher Scientific A21449 | 1:1000 |
| Alexafluor™ Goat anti-rabbit 488 | ThermoFisher Scientific A11034 | 1:1000 |
| Alexafluor™ Goat anti-rabbit 555 | ThermoFisher Scientific A21428 | 1:1000 |
| Alexafluor™ Goat anti-rabbit 647 | ThermoFisher Scientific A21245 | 1:1000 |
| Alexafluor™ Goat anti-rat 555 | ThermoFisher Scientific A21434 | 1:1000 |
| Alexafluor™ Goat anti-mouse 488 | ThermoFisher Scientific A11029 | 1:1000 |
| Alexafluor™ Goat anti-mouse 555 | ThermoFisher Scientific A21424 | 1:1000 |
| Alexafluor™ Goat anti-mouse 647 | ThermoFisher Scientific A21236 | 1:1000 |
| Alexafluor™ Donkey anti-mouse 555 | ThermoFisher Scientific A23424 | 1:1000 |
| Alexafluor™ Donkey anti-mouse 647 | ThermoFisher Scientific A31571 | 1:1000 |
| Alexafluor™ Donkey anti-rabbit 555 | ThermoFisher Scientific A31572 | 1:1000 |
| Alexafluor™ Donkey anti-goat 647 | ThermoFisher Scientific A21447 | 1:1000 |
| Alexafluor™ 488 conjugated affiniPure donkey anti-chicken | Jackson labs 703-545-155 | 1:250 |
| Rhodamine red-X-conjugated affiniPure donkey anti-rat IgG | Jackson labs 712-295-153 | 1:250 |

Images were acquired on a Zeiss fluorescence microscope (Zeiss Airyscan Confocal), using 20x, 40x and 63x oil magnifications and analyzed using Zeiss Zen software. Three to five images were obtained for each compartment of each microfluidic device, and 3 microfluidic devices were utilized for each condition. Cell counts were performed by an individual blinded to the experimental conditions (19).

### Quantitative analysis of neurite outgrowth

For the morphological analysis of neurons, we used TUJ1 immunostaining and 40x oil magnification images. We first traced the neuronal somatic area manually and tracked individual neurites using the simple neurite tracer software plugin on the Fiji package of ImageJ (75). We then performed a Sholl analysis on the neurite mask, with ten soma-centered concentric circles of increasing radii (10µm increment) (20). For analysis, 30 neurons were randomly selected per condition, and we calculated the number of intersections/radius and the total sum of intersections for each individual neuron.

### Trans-synaptic tracing with Rabies Virus

hiPSC-derived spinal neurons were transduced with a G-deleted rabies virus expressing Cre recombinase and enhanced green fluorescent protein (EGFP) (Rabies GΔ Cre-EGFP, Salk Institute, La Jolla, California) at DIV50. After a week, the spinal neurons were co-transduced with the helper virus AAV-CAG-FLEX-oG-WPRE-SV40-PA (Salk Institute, La Jolla, California), which expresses the rabies virus glycoprotein (RVG) required for retrograde trans-synaptic transfer of Rabies-GFP from the spinal neurons to cortical neurons.

For the cortical compartment, cortical neurons were transduced with an AAV9 vector containing a DIO construct for mCherry expression under the control of the CAMKII promoter (AAV9 CAMKII-DIO-mCherry) at DIV44 when seeded at the device to optimize transduction rate. Expression of mCherry was induced by Cre recombinase, which was provided by the Rabies GΔ Cre-EGFP virus, confirming trans-synaptic communication. Confocal fluorescence microscopy (oil immersion 40x) was used to detect both EGFP and mCherry signals in the cortical compartment.

### MEA electrophysiological recordings

Electrophysiological recordings were conducted using the Maestro Edge system (Axion BioSystems) with bonded microfluidic devices placed on 64-electrode 6-well plates. In each well, 16 to 20 electrodes were located beneath or between the microchannels, where signal artifacts arose due to artificial amplification or due to lack of cell coverage at this part of the surface. These electrodes were excluded from analysis. Plates were pre-coated with Geltrex prior to cell seeding, and electrophysiological parameters were recorded as previously described (74).

We conducted five-minute weekly recordings of spontaneous activity for a period of 12 weeks. Both basic and synchronous electrophysiological metrics were examined. Voltage measurements were filtered using the neuronal spontaneous activity filter of the software AxisNavigator. Spikes were identified as instantaneous time points of voltages that exceed 5 spikes/sec. In addition to filtering, all traces were visually inspected for quality control. Hypersynchronous (or “network”) bursts were defined using the envelope algorithm of the software (Neural Metrics Tool, Axion Biosystems software). To ensure proper electrical connectivity and eliminate recording artifacts caused by the placement of the microfluidic device, the wells were flooded with culture medium to establish a fluidic/electrolyte bridge between the interior and exterior of the device, facilitating communication with the ground electrode. Results were graphed using NeuralMetricTool that generates raster plots of the 300sec spontaneous recording. Graphs were generated to depict key electrophysiological metrics over the 12-week experimental period, including weighted mean firing rate (wMFR), number of active electrodes, burst percentage, average burst frequency, network burst frequency, burst duration, synchrony index, and area under the normalized cross-correlation (AUNCC).

For optical stimulation studies, neurotransmitter modulators were tested by applying the AMPA/kainate receptor antagonist NBQX (2,3-dioxo-6-nitro-7-sulfamoyl-benzo[f]quinoxaline, 50uM, Sigma Aldrich, St. Louis, MO, USA) and the NMDA receptor antagonist APV (2-Amino-5-phosphonovaleric acid, 28.4uM). Neurons expressing channel rhodopsin were activated with blue light with wavelength center of 475nm and a maximum intensity of >3.9mW/mm^3^. Voltage measurements were filtered using the optical stimulation filter in AxisNavigator. Optogenetic stimulation was performed using a modified Axion BioSystems setup, allowing light delivery into the 6-well plates containing the PDMS microfluidic devices. Recordings were 5-minute long, with optical stimulation delivered as 5-millisecond (blue or far-red) light pulses at 30% intensity, 0.25 Hz, for a total of 70 stimuli. Events spanning 100 milliseconds of peristimulus time with a bin size of 1 millisecond were analyzed and visualized using Neural Metric Tool (Axion Biosystems).

### Statistical Analysis

All data were analyzed using Graph Pad Prism software 10.4.1 (Boston, MA, USA). Data are presented as mean ± SEM. For experiments comparing two conditions over time, we used repeated measures two-way ANOVA when assumptions of normality and equal variances were met. For datasets with small sample sizes or non-normal distributions, we used Wilcoxon signed-rank test on values to assess directional significance. Each experiment included at least three technical replicates. Experimental conditions and technical replicates are specified in the figure legends. Statistical significance for all tests performed was set at *P*<0.05.

## Acknowledgements and funding sources

This manuscript was supported by National Institute of Health 5R01NS117604 and Maryland Stem Cell Research Funding grant 2023-MSCRFD-6125 (NJM), National Institute of Health R25NS065729 (AT). National Institute of Health K08NS102526, Doris Duke Foundation Clinical Scientist Award to CWH.

