## Supplemental Figures 1-2 for "Development of a Human iPSC-Derived “Corticospinal Tract-on-a-Chip” for Neurodegenerative Disease Research"

Supplemental 1

**A** *Addition of live cell dye at the cortical compartment only*

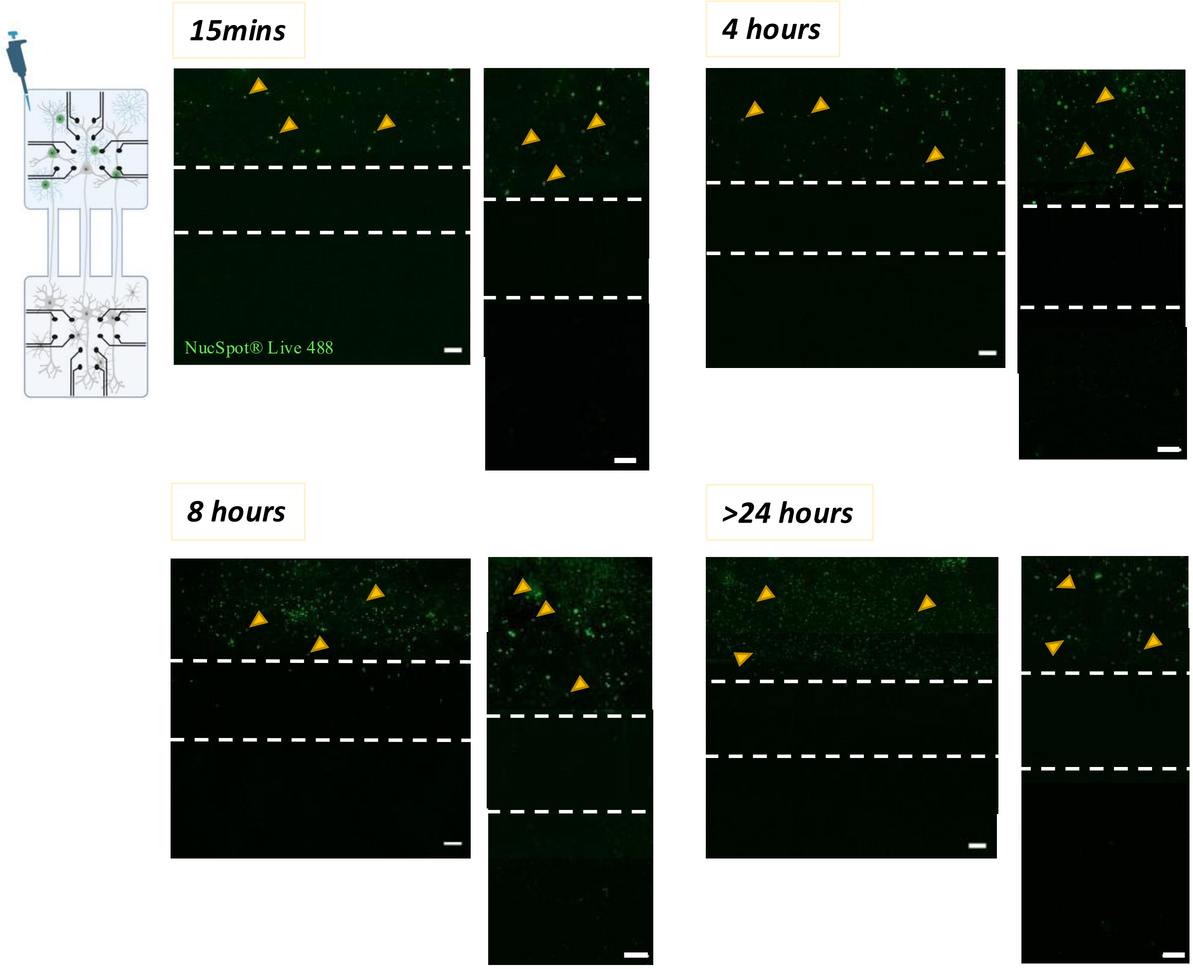


**B** *Subsequent addition at the spinal cord compartment*

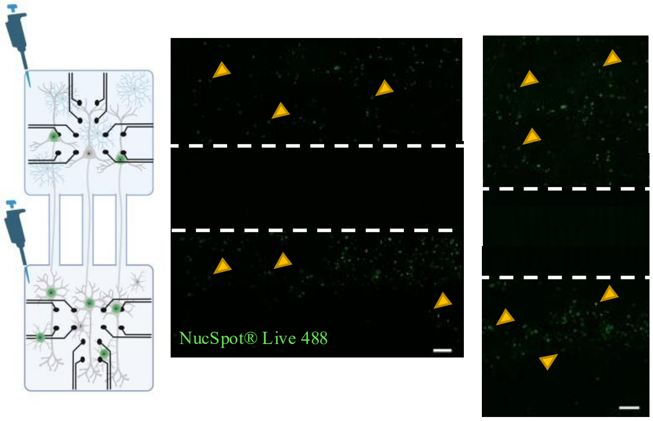

**Supplemental 1: Live cell diffusion demonstrates fluidic isolation between compartments.**

**A.** Time-lapse imaging of dye diffusion in the microfluidic device following the addition of a live-cell dye to the cortical compartment. Mosaic assembled manually from overlapping confocal fields to represent the broader field of view. Images were captured at 15 minutes, 4 hours, 8 hours, and >24 hours post-addition to monitor diffusion dynamics across the microchannels (n=3 per condition). **B.** Representative mosaic live-cell images of microfluidic devices after dye addition to both compartments, demonstrating expected diffusion patterns (n=3). Scale bars=100μm.

### Supplemental 2

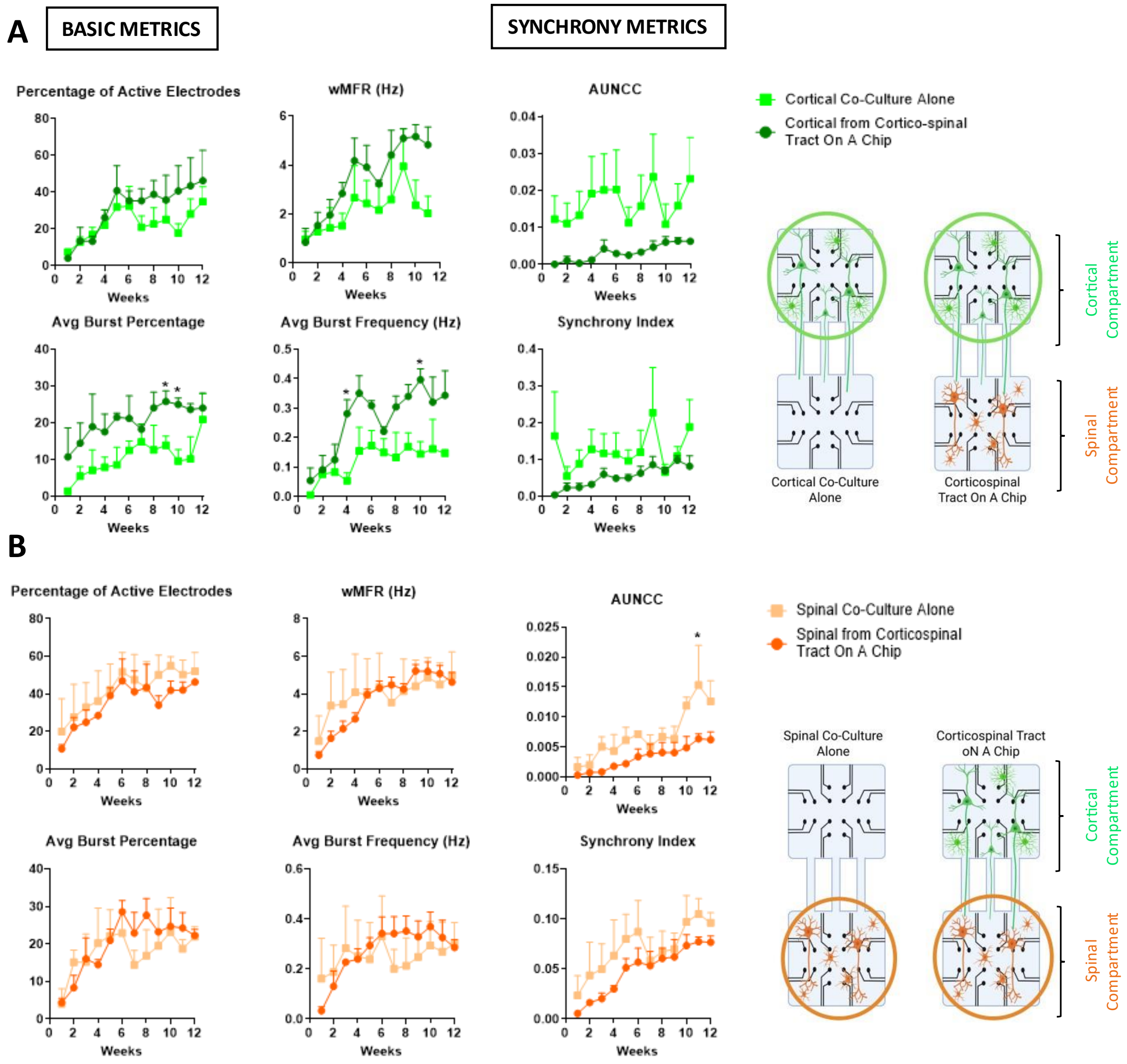

**Supplemental 2: Comparison of electrophysiological activity and maturation of single population chips (one neuronal population) with CST-on-a Chip (two neuronal populations)**

**A.** CNs and CA single population cultures compared to CST-On-A-Chip setup with both cortical and spinal co-cultures **B.** SpMN and SpA compared to CST-On-A-Chip setup, and illustration of both plating setups (n=8-9 devices on MEA per condition). One-way Anova, * $p<0.05$, wMFR: weighted mean firing rate, AUNCC: area under normalized cross correlation.
